# NGS-enriched activated sludge modelling of industrial wastewater treatment plant

**DOI:** 10.1101/2023.01.23.523537

**Authors:** M A Prawira Negara, K J Keesman, G J W Euverink, B Jayawardhana

**Author notes:** (M.A.P. Negara); (K.J. Keesman); (G.J.W. Euverink); (B. Jayawardhana).

## Abstract

Metagenomics advances with the Next Generation Sequencing (NGS) technology offer detailed insights into the microbial communities and their activities in a Wastewater Treatment Plant (WWTP). Since it has been shown recently that the microbial communities can be related to process data through machine learning, we investigate in this paper the enrichment of Activated Sludge Model 1 (ASM1) using time-series NGS data. We first present a modified ASM1 (mod-ASM1) to describe the industrial wastewater treatment at North Water’s WWTP facility in Delfzijl, the Netherlands. Subsequently, we identify the parameters for the ten weeks (weeks 40-50, 2014) of process data from North Water WWTP with prior parameters from the recommended ones from IWA. We further established a subset of parameters that are correlated to NGS data. Based on this relationship, a parameter-varying mod-ASM1 is obtained where the parameter variation is directly linked to the NGS data. We validate the NGS-enriched mod-ASM1 in the prediction of process data in the subsequent three weeks (weeks 50-53, 2014). While the enriched mod-ASM1 gives a good estimation of the COD effluent data, it cannot capture the production of nitrogen, which is often missed when the static model is deployed.

## 1. Introduction

Wastewater forms a significant part of waste from human activities, and it requires treatment before it is environmentally safe to be discharged into the natural water resources (see, for example, Henze and Comeau (2008)). When it is discharged untreated, it pollutes water resources and can lead to disastrous ecological, environmental, as well as economic impacts. Wastewater comes from two major sources: human sewage systems and process waste from manufacturing industries. The biological treatment of wastewater was first introduced in the early twentieth century and has become the basis of wastewater treatment worldwide. Activated sludge, which is composed of a high concentration of microbial communities of bacteria, protozoa, and archaea, is used in this biological treatment. These microbial communities consume organic compounds, ammonia, and phosphate for their activities and growth and thus cleanse the wastewater. After the clarification step (where the activated sludge is sedimented), the treated wastewater can be safely discharged to natural water resources, such as rivers or seas.

Although the treatment process operations appear to be straightforward, the underlying biological-chemical process is complex (Davies (2005)) and difficult to monitor. Consequently, the process control relies mainly only on the physical process variables that can be measured, and there is no direct monitoring and control over the microorganism activities. It is well-known that the dynamical variation in WWTP is due to the time-varying composition and activity of the microorganism in the reactors affected by the variation of influent nutrient composition. The influent can vary in the flow rate, chemical composition, pH, and temperature, which in turn affect the population dynamics and metabolic process of the activated sludge. In some cases, the dynamical variations are also affected by the return sludge. Notably, some WWTPs that treat industrial wastewater, as the subject of this paper, must deal with recalcitrant chemicals that are hard to degrade by the activated sludge. Such industrial wastewater contains harmful chemicals and has extreme properties such as low/high pH or high salt concentration, which directly affects the performance of the activated sludge.

There are a number of established models for describing the process dynamics in WWTP, such as the well-studied Activated Sludge Models from the International Water Association (see, for instance, Henze et al. (2000)). These models have been shown to be effective in establishing dynamical relationships between the activated sludge and the decomposition of organic and inorganic matter from the influent (see Wang et al. (2017)). In these models, the competition of the microbial community affects the parameters in the kinetics of the WWTP processes such as maximum specific growth rates and affinity constants. These parameters represent the combined metabolic activities of a group of microorganisms and can therefore have a large degree of variations depending on the growth conditions of the microorganisms in the activated sludge. Due to this uncertainty, the models have not been used for real-time model-based optimization and control of WWTP. Instead, WWTP is assumed to always operate in a quasi-steady-state condition, and process control adjustments are done manually and in an *ad hoc* fashion based on the perceived changes in the process parameters.

For improving the reliability and applicability of these models, it has recently been acknowledged that the knowledge and real-time information of the microorganism population are very important to enrich the ASMs in real-time, as concluded in Muszynski et al. (2015), Liu et al. (2016) and Bassin et al. (2017). The microbial community composition is very dynamic, which usually leads to changes in the metabolic functional capabilities of the community and reflects the underlying biotic and abiotic processes (Werner et al. (2011), Kim et al. (2013), Hai et al. (2014)). Therefore understanding the microbial temporal pattern of activated sludge will provide insight into the role of different microorganisms and allow us to optimize the treatment process performance (Briones and Raskin (2003)). For describing the population dynamics of the microbial community in WWTP (without relating them to the metabolic functions), a number of models have been developed. These include the neutral community model (see Sloan et al. (2006) and Ofiteru et al. (2010)), the mass balance model (see Saunders et al. (2016) and Mei et al. (2016)), the mass-flow immigration model (see Frigon and Wells (2019) and Guo (2019)) and the non-steady-state mass balance model (see Sun et al. (2021)).

In this paper, we study the integration of real-time microbial community information obtained from DNA using the Next Generation Sequencing (NGS)^1^ technology into the ASMs. Firstly, we present a modified ASM (mod-ASM) that closely describes the industrial wastewater treatment plant where organic material is largely present in the influent. Based on the mod-ASM, we fit some of the parameters based on the measured process data using prior information of parameters from IWA (Henze and Comeau (2008)). A few parameters are coupled to the microbial community through nonlinear models based on the established correlation of the genera to the state variables as reported in Negara et al. (2019). This method allows us to obtain a parameter-varying ASM that couples the microbial community dynamics from NGS to the ASM.

The rest of the paper is organized as follows. In Section 2, we explain the materials, data, and methodology that we use in this study. In Section 3, we present the development of NGS-enriched mod-ASM1. In Section 4, numerical results are presented where the model is fitted to the process, and NGS data is validated and used as a predictive model. The paper is concluded in Section 5.

## 2. Materials and methods

### 2.1. North Water’s wastewater treatment plant

For the material and data used in this study, we used the process data and NGS data that were taken from North Water’s Saline Wastewater Treatment Plant (SWWTP) in Delfzijl, the Netherlands. This industrial SWWTP^2^ treats collective industrial wastewater from chemical industries that are operated in a chemical park in Delfzijl and its neighborhood. The processed clean water is then released into the surface water. General operational data of this SWWTP are given in Table 1.

**Table 1.**
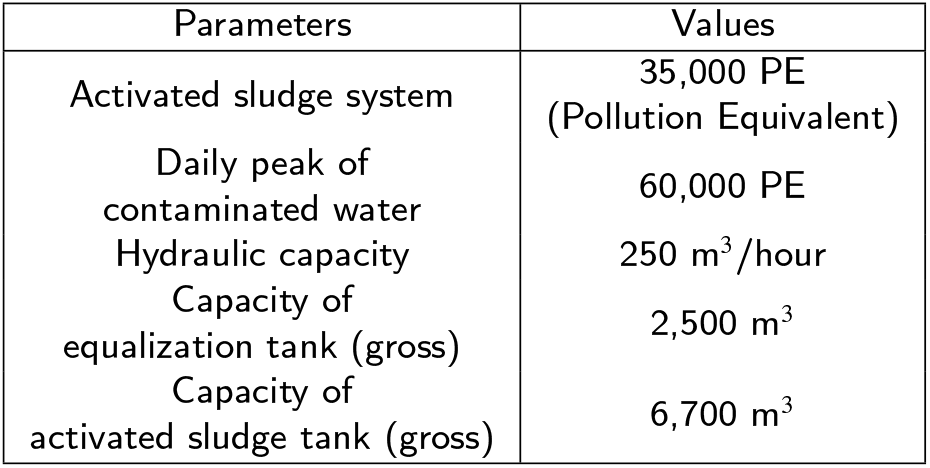
North Water’s SWWTP Purification Data

| Parameters | Values |
| --- | --- |
| Activated sludge system | 35,000 PE<br>(Pollution Equivalent) |
| Daily peak of contaminated water | 60,000 PE |
| Hydraulic capacity | 250 m<sup>3</sup>/hour |
| Capacity of equalization tank (gross) | 2,500 m<sup>3</sup> |
| Capacity of activated sludge tank (gross) | 6,700 m<sup>3</sup> |

A general schematic of the biological treatment process of North Water’s SWWTP is shown in Fig. 1. As shown in Fig. 1, the influent coming from various sources is collected in the equalization tank (blending) before it is pumped into the activated sludge tank. The anaerobic biological process is first taken place in the inner part of the activated sludge tank before proceeding to the aerobic process in the outer part of the activated sludge tank. In the clarifier tank, the activated sludge is separated from the water and is partly fed back to the activated sludge tank while the treated water is then discharged into the Wadden Sea.

**Figure 1:**
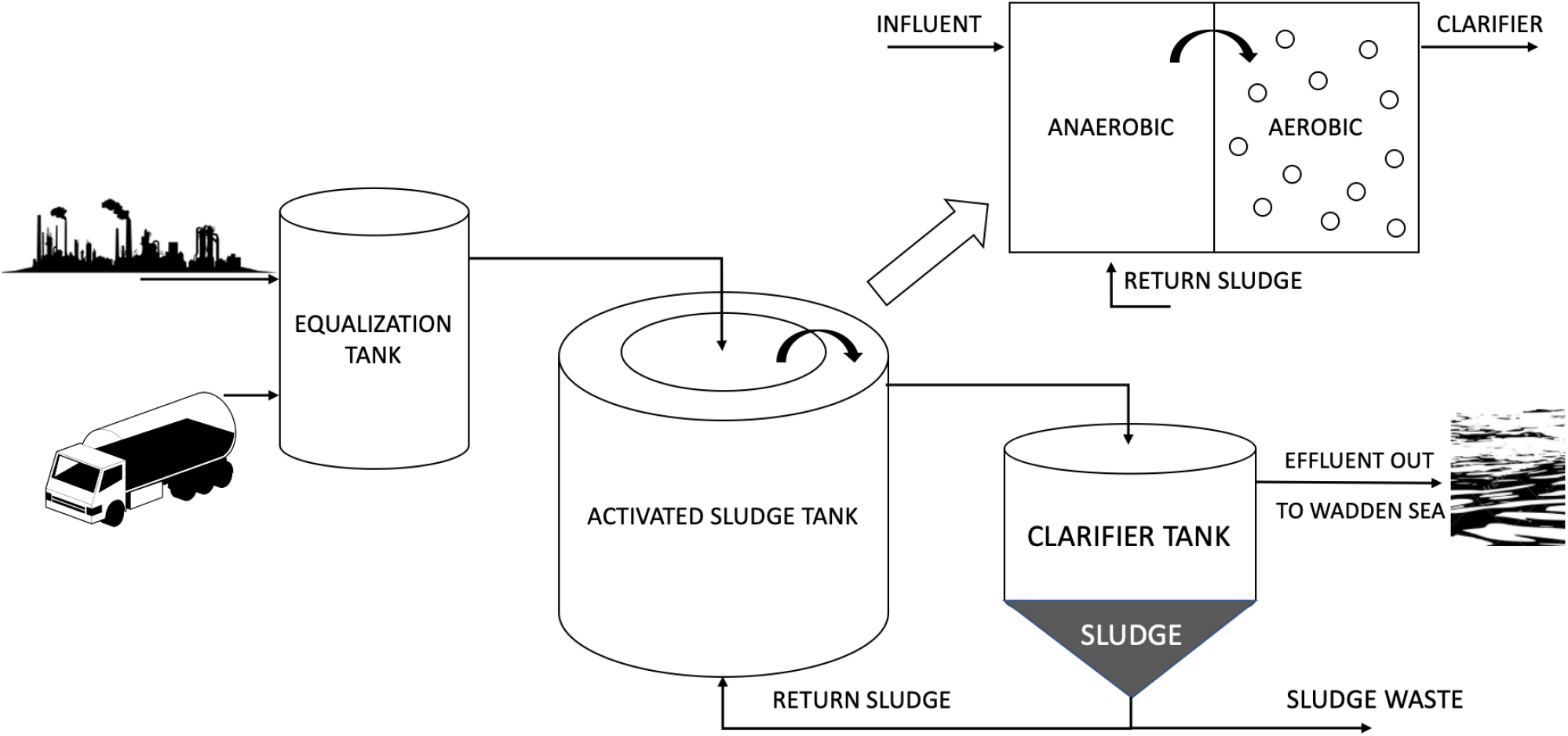
Schematic diagram of the North Water’s Saline Wastewater Treatment Plant, where the influent from the industry is processed and the effluent is discharged into the Wadden Sea.

From Fig. 1 we can see that the process in the activated sludge tank can be regarded as a two-reactor system of anaerobic and aerobic processes, respectively, that are connected sequentially. The setup of an anaerobic process followed by an aerobic one is atypical for standard WWTP. This reverse setup is possible for North Water’s WWTP since the influent has a high chemical oxygen demand (COD) such that it contains sufficient energy in the wastewater for the removal of nitrite and nitrate in the first reactor. The complete aeration process takes place in the aerobic reactor part, where industrial waste is further broken down by biological processes and consumes the rest of the ammonium.

We collected the process data from this SWWTP and used it in our simulation model. The data were collected from week 40 in 2014 until week 6 in 2017 on an irregular basis and processed as a weekly dataset. For fitting our simulation model to the data later, we use Piecewise Cubic Hermite Interpolating Polynomial (PCHIP) to obtain daily measurement data.

### 2.2. Activated Sludge Models

As described in the Introduction, a number of Activated Sludge Models (ASM) have been used in literature to mathematically describe the kinetics of biological processes in WWTP. Some of the established models from the International Water Association are the ASM1, ASM2, ASM2d, and ASM3, and we refer interested readers to the exposition in Henze et al. (2000). These models incorporate various biological activities of the active sludge that is dominated by bacteria and protozoa. Common biological processes that are included in these models are the oxidation of carbonaceous biological matter, the oxidation of nitrogenous biological matter (e.g., ammonium and nitrite), and removing nutrients (mainly nitrogen and phosphorus).

In this paper, we focus on the use of ASM1 to model North Water’s SWWTP since this model contains most of the important biological processes. In particular, ASM1 contains nitrogen and COD removal processes within the suspended-growth treatment process, as well as the nitrification and denitrification processes, as discussed in Nelson and Sidhu (2009). These are the main processes that take place in the activated sludge tank of North Water SWWTP as shown in Fig. 1. In this SWWTP, 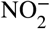 and 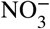 are removed in the first tank, while COD and 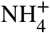 are mostly removed in the aerated second tank.

Based on ASM1 as our prototype model, we present a modification of ASM1 to model the sequentially operated anaerobic and aerobic phases in the activated sludge tank. We recall that the standard ASM1 uses 13 state variables and 8 process kinetics. By denoting *S* as the soluble matter and *X* as the particulate ones, the state variables in ASM1 are the inert organic matters (*S_I_*&*X_I_*), biodegradable substrates (*S_S_&X_S_*), active biomass (*X_BA_&X_BH_*), particulate product from biomass decay (*X_P_*), oxygen (*S_O_*), nitrite and nitrate (*S_NO_*), ammonium and ammonia (*S_NH_*), biodegradable organic nitrogen (*S_ND_&X_ND_*), and alkalinity (*S_ALK_*). The kinetic processes in ASM1 are aerobic growth of heterotrophs and autotrophs, anoxic growth of heterotrophs, decay of heterotrophs and autotrophs, ammonification, hydrolysis of entrapped organic and entrapped organic nitrogen.

Since North Water’s SWWTP processes industrial wastewater, the influent contains a small amount of nitrogen which is also observed in the very low measurement levels of *X_P_*, *S_ND_*, *X_ND_* and *S_ALK_*. These four state variables also do not have any direct impact on the process kinetics of ASM1. In addition to potentially removing these four state variables from the model, the inert organic matters of *S_I_* and *X_I_* can also be removed since they are external variables in the original ASM1 and do not play a role in the dynamics of SWWTP. Therefore we modify the standard ASM1, where we reduce the number of state variables from 13 to 7 states. Consequently, the number of kinetic processes is also reduced from 8 to 7, where hydrolysis of entrapped organic nitrogen is removed from the reduced model. The mod-ASM1 for describing North Water’s SWWTP is summarized in Table 2 containing all process kinetics *ρ*(*x*), state variables *x_i_*, and the stoichiometric matrix *w*. Correspondingly, based on the work of Benhalla et al. (2010), the mod-ASM1 for North Water’s SWWTP is given by

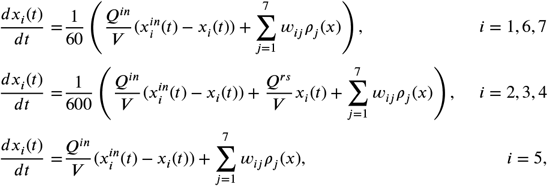

where *Q^in^* is the flow rate of the influent, *Q^rs^* is the flow rate of the return sludge, *V* is the total volume of the reactor, *w_ij_* is the stoichiometric constant for the state variable *x_i_* in the *j*-th reaction, and *ρ* is the process kinetics as shown in Table 2. Note that in this model, the time *t* is in seconds, and the constants 1/60 and 1/600 reflect the time-scale separation between the different processes that are used in the simulation. The time-scale constants above are chosen based on the approximate time constant for the different processes and on the simulation performance.

**Table 2.**
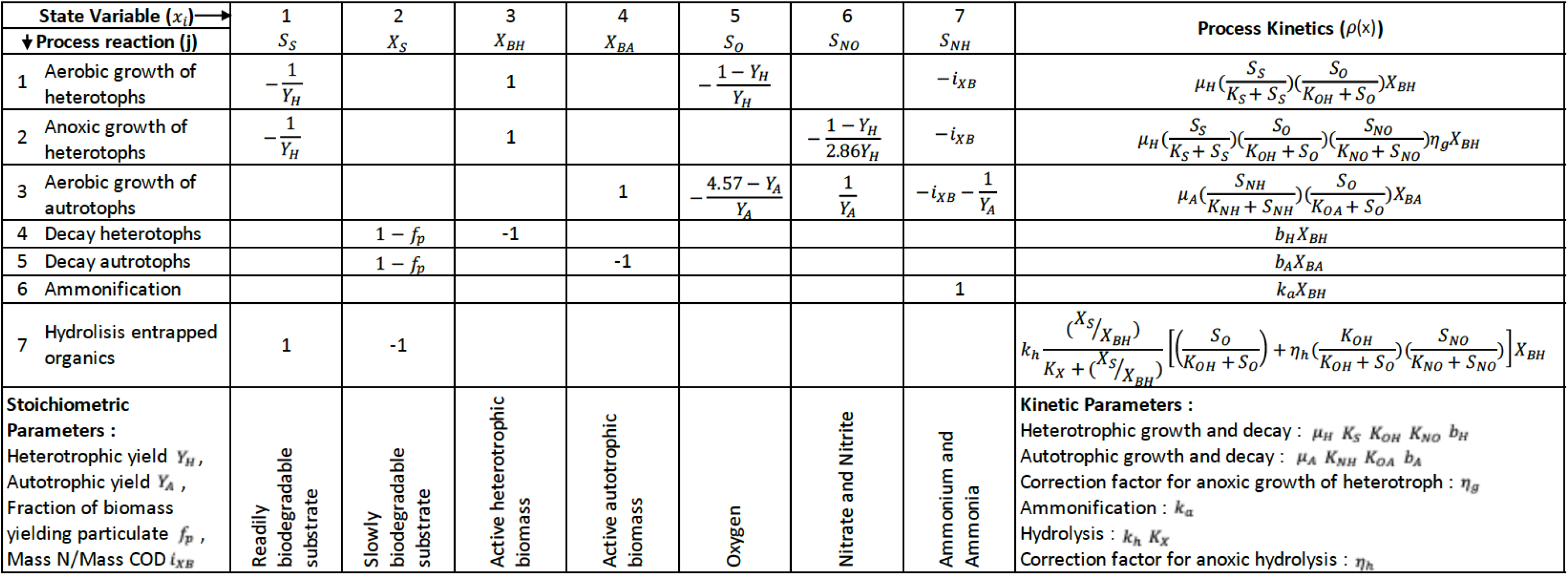
The stoichiometric matrix *w* in the modified ASM1 that relates 7 reaction rates *ρ_j_*(*x*) to the 7 state variables *x_i_*. The reaction rates/process kinetics is given in the rightmost column, while the description of parameters & state variables is given at the bottom of the table.

### 2.3. NGS data analysis

The NGS dataset, taken from the WWTP in Oosterhorn, was made available for this paper by Bioclear earth. The sample data were taken from week 40 in 2014 until week 10 in 2017. The dataset was arranged in multiple sheets of data, each sheet representing one hierarchical rank of taxonomy. The available taxonomical ranks in the dataset are (in hierarchical order): Class, Order, Family, and Genus. For this study, the genus rank was chosen to develop and build the models. This rank shows the highest level of detail of the four available ranks. In total, 1236 different genera were identified in the 32 samples.

In the dataset, samples are taken on average every four weeks, resulting in 32 total samples. Due to practical constraints, these samples were not always taken over a regular four-week period. The sequencing results were then normalized to fractions. The normalized fraction was calculated as log *x*, where *x* is the value of NGS-reads for a specific order divided by the total value of NGS-reads. Since the process data has a higher weekly frequency, the NGS dataset is re-sampled into a weekly frequency using interpolation. In this paper, we apply the Piecewise Cubic Hermite Interpolating Polynomial (PCHIP) method as it can approximate the missing data points while satisfying the positive definite constraint. Figure 2 shows the time-varying data of some of the microorganisms’ population dynamics with the interpolated dataset using PCHIP method.

**Figure 2:**
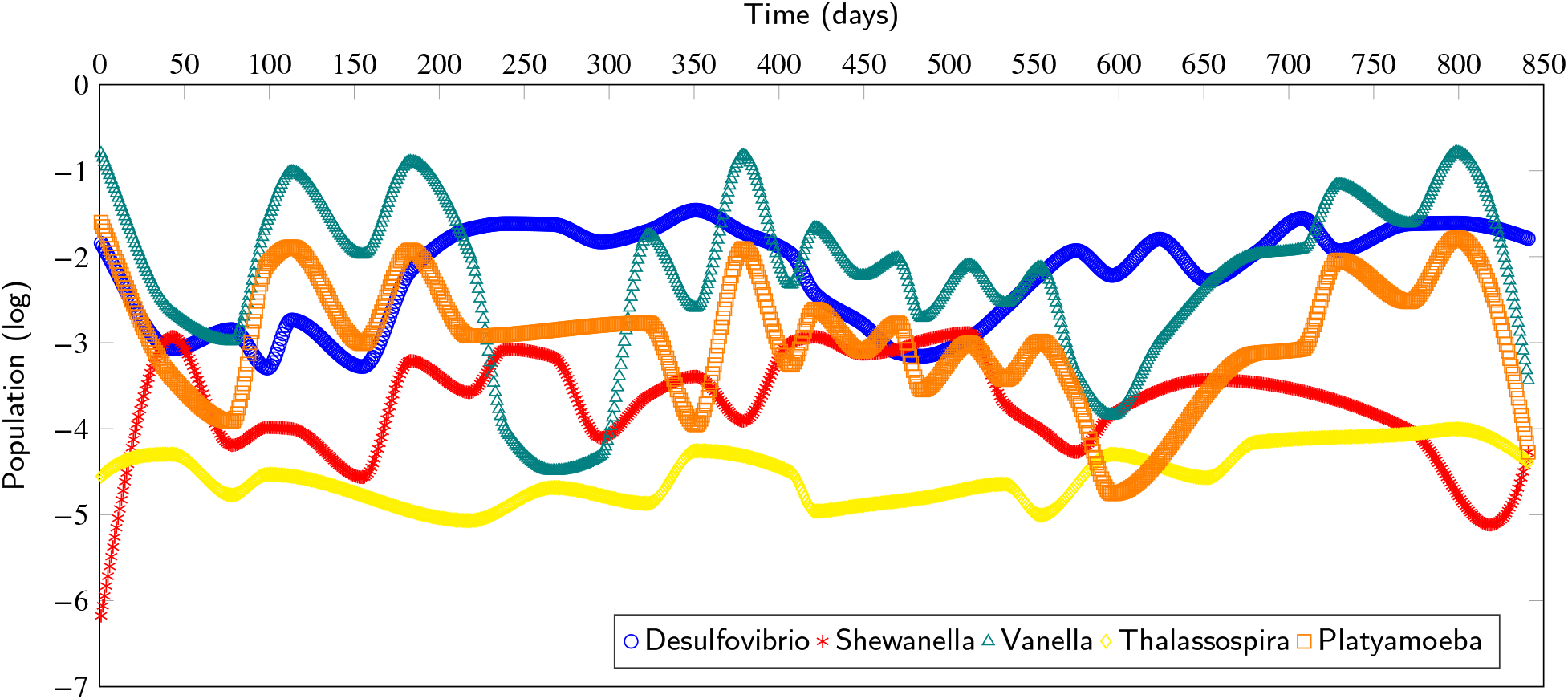
The microbial population time-series of five genera based on the NGS analysis of samples from North Water’s SWTTP collected from week 40 in 2014 until week 6 in 2017 with an interval of around four weeks. The PCHIP interpolation method is used to obtain the daily data.

The analysis we use for this dataset is based on a machine learning technique, the so-called Support Vector Regression (SVR), which can handle well high-degree of uncertainties and unstructured relations between rich sets (high degree of heterogeneity) of input and output data (with a small number of time points). Such preliminary analysis can be extremely useful in identifying a subset of the microbial population that can explain most of the variations observed in the process dynamics.

In Negara et al. (2019) by analyzing the (local) sensitivity of each modeled process variable to each genus (NGS data) and based on the SVR found in the previous step, an indication of the influence of the microbial structure on process performance was found. Table 3 shows the sensitivity analysis (SA) of four process variables related to the genera found in the system. One Factor At a Time (OFAT) technique is used for this SA. According to Razavi and Gupta (2015) OFAT is a local sensitivity analysis technique that computes a finite-difference approximation of the local slope of the response surface around a base point in the factor space. In this case, the approximation determines the influence of a genus on process variables for a certain change of that genus. Thus, only a local area of the influence of each genus is explored.

**Table 3.**
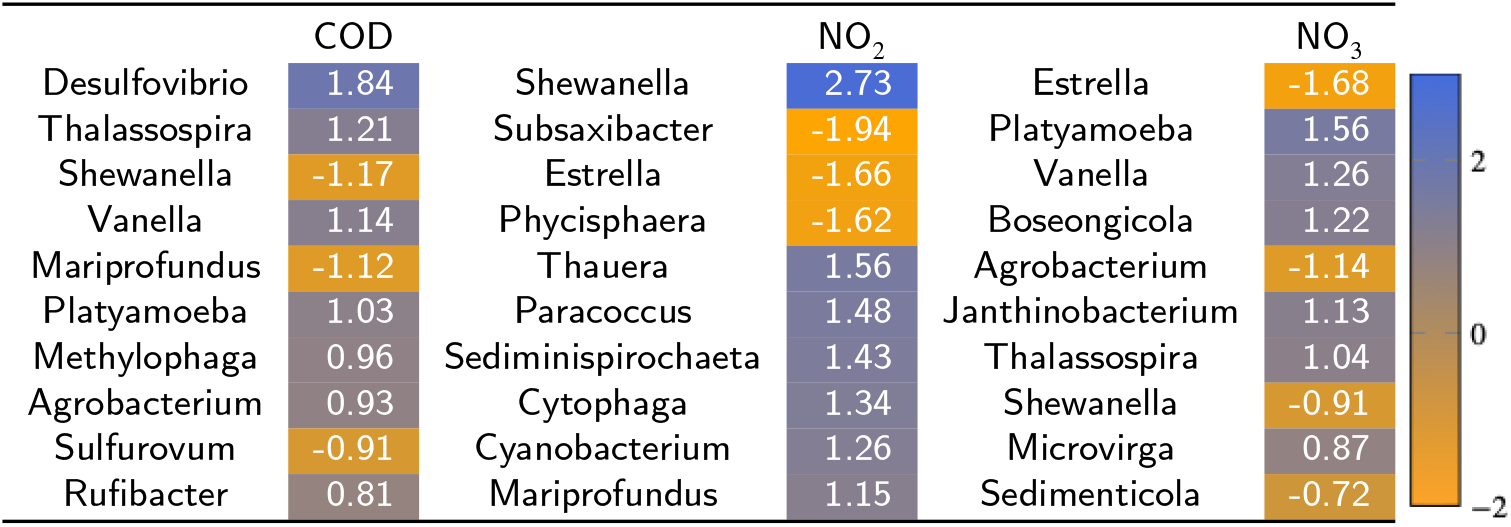
Sensitivity analysis of the top 10 genera using OFAT on the trained model SVR for different process components where the positive value signifies positive relation and vice versa.

## 3. Coarse-grained modeling

Coarse-grained modeling is a method of modeling that aims at simulating the behavior of complex systems using their coarse-grained (simplified) representation. In this study, we simplify the heterogeneity of microorganisms found in the wastewater treatment plant through the explained method in the previous section and relate it to some of the mod-ASM1 parameters. Since the system that we want to model has two reactors that work sequentially where the second reactor influent comes from the first reactor effluent, we describe the system with two mod-ASM1. Both models are assumed to have the same set of parameters for the two reactors where the effluent of the first mod-ASM1 becomes the influent of the second mod-ASM1.

The mod-ASM1 we use describes a continuous system that is fed with influent on a daily basis. Because the data we have is a weekly data set, we perform the same PHCIP interpolation technique to obtain a daily data set that will be used in our simulations using the coarse-grained mod-ASM1. We note that the application of linear systems identification to the data does not yield a good fit, and it can even lead to unstable linear systems. This shows that the underlying dynamical systems are non-linear (as given in the well-known ASM and in our mod-ASM1). For parameter estimation and model validation, we use the data from the first 100 days (week 40 in 2014 until week 5 in 2015). Particularly, the dataset from the first 70 days is used to identify the coarse-grained mod-ASM1 whose parameters are coupled to the microbial population from the NGS dataset, and subsequently, the subsequent 30 days dataset is used to show the predictive capability of the NGS-enriched mod-ASM1.

### 3.1. Process parameters estimation

Before we couple the microbial community data with the mod-ASM1, we perform a two-step parameter estimation of the mod-ASM1 based on the process data. In the first step, we performed a Monte-Carlo method for parameter estimation, where 40 different sets of measurement data are sampled according to the expected measurement noise in the process data. The dataset can be seen in Appendix A. This procedure allows us to obtain the possible time-variation of the parameters within a range that can later be linked to the microbial community data. The parameter estimation is performed by finding parameters that minimize a quadratic cost function given by the sum-of-square of the relative errors between the model output and measured process data. As reported in the literature, see, for instance, Abusam et al. (2001), Wang et al. (2017), Muoio et al. (2019), Du et al. (2020), Vitanza et al. (2021) and Wu et al. (2021), the fitted parameters can describe the different characteristics of different wastewater treatment plants. In the second step, we hypothesize that the time-variation of these parameters is caused by the changes in the biological process in the reactors, which are reflected by the dynamics in the biological composition of the activated sludge. The information on the latter can be provided by the NGS data.

In the mod-ASM1, there are five stoichiometric parameters of biochemical reactions and 14 kinetic parameters. The parameter estimation was carried out using the data from week 40 to week 50 in 2014. The process data of effluent COD and 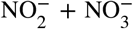 were used to estimate the parameters. Using the sum-of-square of relative errors as the cost function, we performed parameter estimation using a number of solvers in Matlab as given in Table 4. In this parameter estimation step, 17 out of 19 parameters are assigned to be static (constant) parameters, while two affinity parameters (*K_OH_&K_NO_*) are designated as dynamics parameters that will later be coupled to the NGS data. This choice is based on the high degree of correlation between these parameters and the influent data.

**Table 4.** Root Mean Square Error from different estimation methods performed in MATLAB.

| Method | RMSE |
| --- | --- |
| fmincon | 4.2382 |
| fminsearch | 5.0132 |
| lsqcurvefit | 3.9338 |
| lsqnonlin | 3.9462 |
| patternsearch | 4.1483 |
| particleswarm | 4.7236 |

### 3.2. Dynamic process parameters

In this subsection, we will present methods to couple the microbial community real-time data from NGS with the time-varying parameters *K_OH_* and *K_NO_*. From the kinetics of mod-ASM1 as given in Table 2, the parameter *K_OH_* is mainly associated with the dynamics of COD (given by the sum of the state variables *S_S_* and *X_BH_*) and related to the aerobic and anoxic growth of heterotrophs. The parameter *K_NO_* is related to the dynamics of 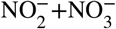 (given by the state variable *S_NO_*). Based on the sensitivity analysis of NGS data and process data using OFAT as presented in Subsection 2.3, we will couple these two parameters and the microbial community through a polynomial relationship as follows

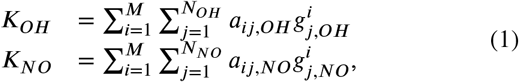

where *a_ij,OH_* and *a_ij,NO_* are parameters of polynomial to be identified, *M* is the degree of polynomial with *M* = 5, *g_j,OH_* and *g_j,NO_* are the log-value of a population of the *j*-th genera out of *N_OH_* and *N_NO_* number of genera that are found to be sensitive to the dynamics of *COD* and of 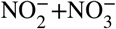, respectively. Using the same method of parameter estimation as before, we estimate the parameters of *a_ij,OH_* and *a_ij,NO_*.

## 4. Results and discussion

Table 5 summarizes the results from the Monte-Carlo simulations along with the baseline parameters from IWA standard (see Henze et al. (2000)). The estimated parameters are presented by the mean value from the Monte Carlo simulations and the corresponding minimum and maximum values. Note that the IWA standard of ASM parameters is mainly based on municipal wastewater. Since in this paper we are dealing with industrial wastewater, which has a different composition of influent from that of municipal one, we do not impose constraints on the parameters with respect to the IWA standard ones during the parameter estimation. This is done in order to remove potential bias in the estimation of the parameters for municipal wastewater.

**Table 5.**
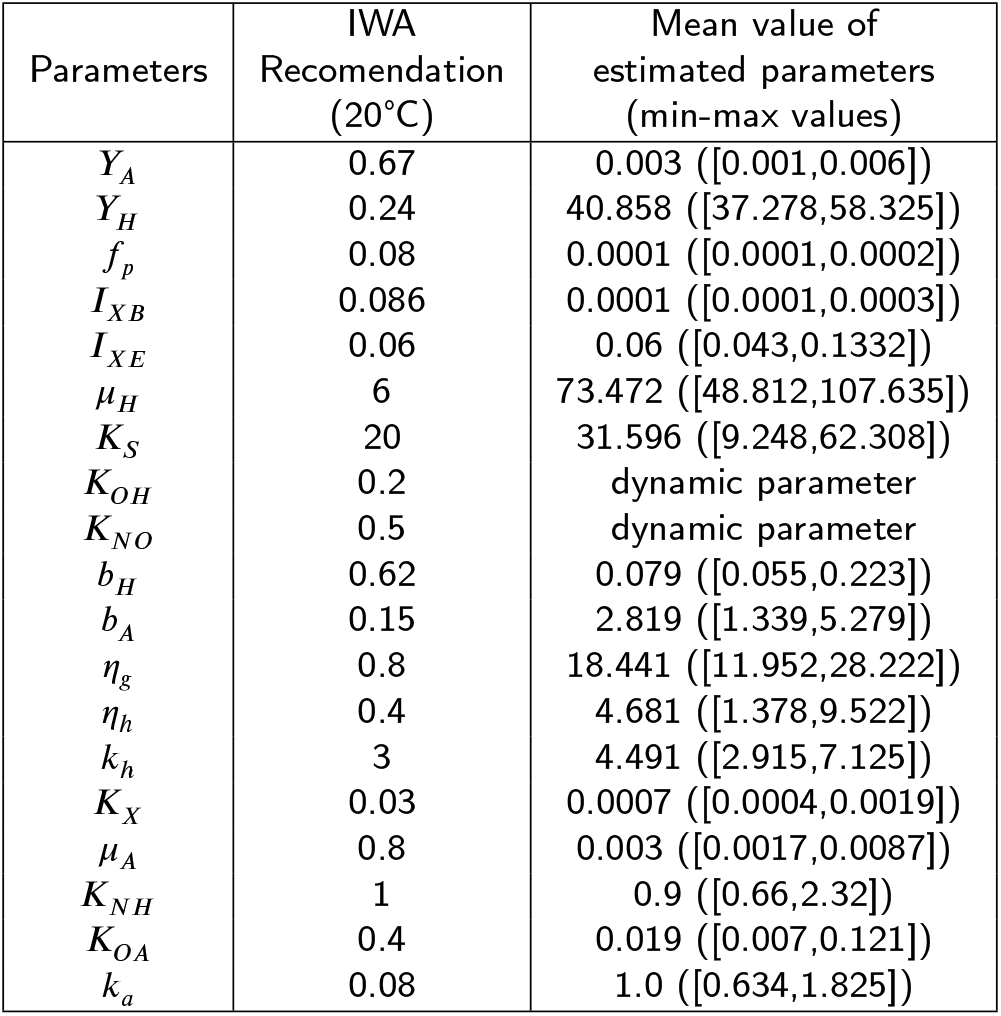
Estimated result of process parameters of ASM1 for North Water’s SWWTP.

| Parameters | IWA Recommendation (20°C) | Mean value of estimated parameters (min-max values) |
| --- | --- | --- |
| $Y_A$ | 0.67 | 0.003 ([0.001,0.006]) |
| $Y_H$ | 0.24 | 40.858 ([37.278,58.325]) |
| $f_p$ | 0.08 | 0.0001 ([0.0001,0.0002]) |
| $I_{XB}$ | 0.086 | 0.0001 ([0.0001,0.0003]) |
| $I_{XE}$ | 0.06 | 0.06 ([0.043,0.1332]) |
| $\mu_H$ | 6 | 73.472 ([48.812,107.635]) |
| $K_S$ | 20 | 31.596 ([9.248,62.308]) |
| $K_{OH}$ | 0.2 | dynamic parameter |
| $K_{NO}$ | 0.5 | dynamic parameter |
| $b_H$ | 0.62 | 0.079 ([0.055,0.223]) |
| $b_A$ | 0.15 | 2.819 ([1.339,5.279]) |
| $\eta_g$ | 0.8 | 18.441 ([11.952,28.222]) |
| $\eta_h$ | 0.4 | 4.681 ([1.378,9.522]) |
| $k_h$ | 3 | 4.491 ([2.915,7.125]) |
| $K_X$ | 0.03 | 0.0007 ([0.0004,0.0019]) |
| $\mu_A$ | 0.8 | 0.003 ([0.0017,0.0087]) |
| $K_{NH}$ | 1 | 0.9 ([0.66,2.32]) |
| $K_{OA}$ | 0.4 | 0.019 ([0.007,0.121]) |
| $k_a$ | 0.08 | 1.0 ([0.634,1.825]) |

Figure 3 shows the plot of WWTP simulations based on the parameters obtained from the Monte Carlo simulations as before. The trajectory in circle is obtained based on the use of mean value parameters as given in Table 5 while the bars show the minimum and maximum trajectories that are obtained from the Monte Carlo simulations. This plot shows that despite a largely good agreement of the WWTP simulations with the process data on the COD, there is still a significant discrepancy in the simulated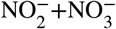. This is due to the fact that two parameters, which play an important role in the dynamics of 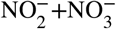, namely *K_OH_* and *K_NO_*, have not been fitted yet to the process data. We also see that there are a couple of misfits that can be seen in the COD from days 20 to 24 and days 49-54. This reading arose possibly because of the use of interpolation in the data, as mentioned in Section 2.1, which made us unable to clearly see if there was a deviation in the influent or effluent at the actual system.

**Figure 3:**
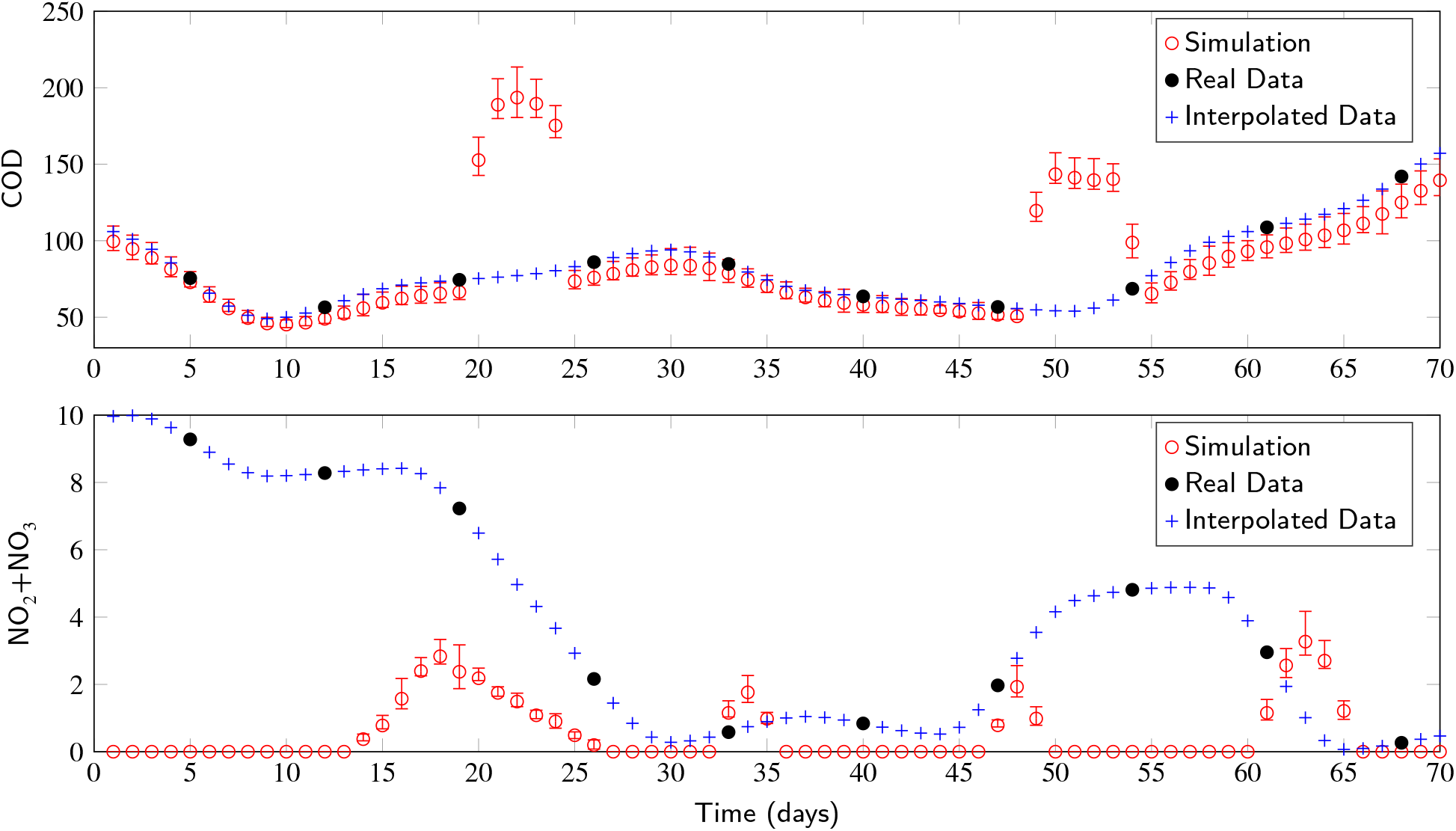
The plot of time-series data of COD and 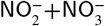 from measurement (shown in blue plusses) and simulations (shown in red circles). The simulation is based on the use of mod-ASM1 where most of the parameters have been fitted to the data except two parameters of *K_OH_* and *K_NO_* that are correlated to the NGS data. In this plot, these two parameters are set to the IWA recommended values as in Table 5. The error bars show the mean value and the min-max values from Monte Carlo simulation.

Using both the static parameters and the dynamics ones (coupled to the NGS data), the resulting simulation and prediction of the process data on COD and on 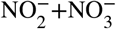 is shown in Figure 4. As shown in this figure, we use the first 70% of the data to train the model according to the methods presented before and the last 30% of the data is used to validate the trained model. The trajectories shown in (dark and light) red crosses are the outcome of the simulation based on fitted parameters with the parameters *K_OH_* and *K_NO_* fixed to the baseline parameters from IWA standard. The trajectories in (light and dark) green circles are obtained using fitted parameters with *K_OH_* and *K_NO_* being coupled to the NGS data according to the identified model in (1).

**Figure 4:**
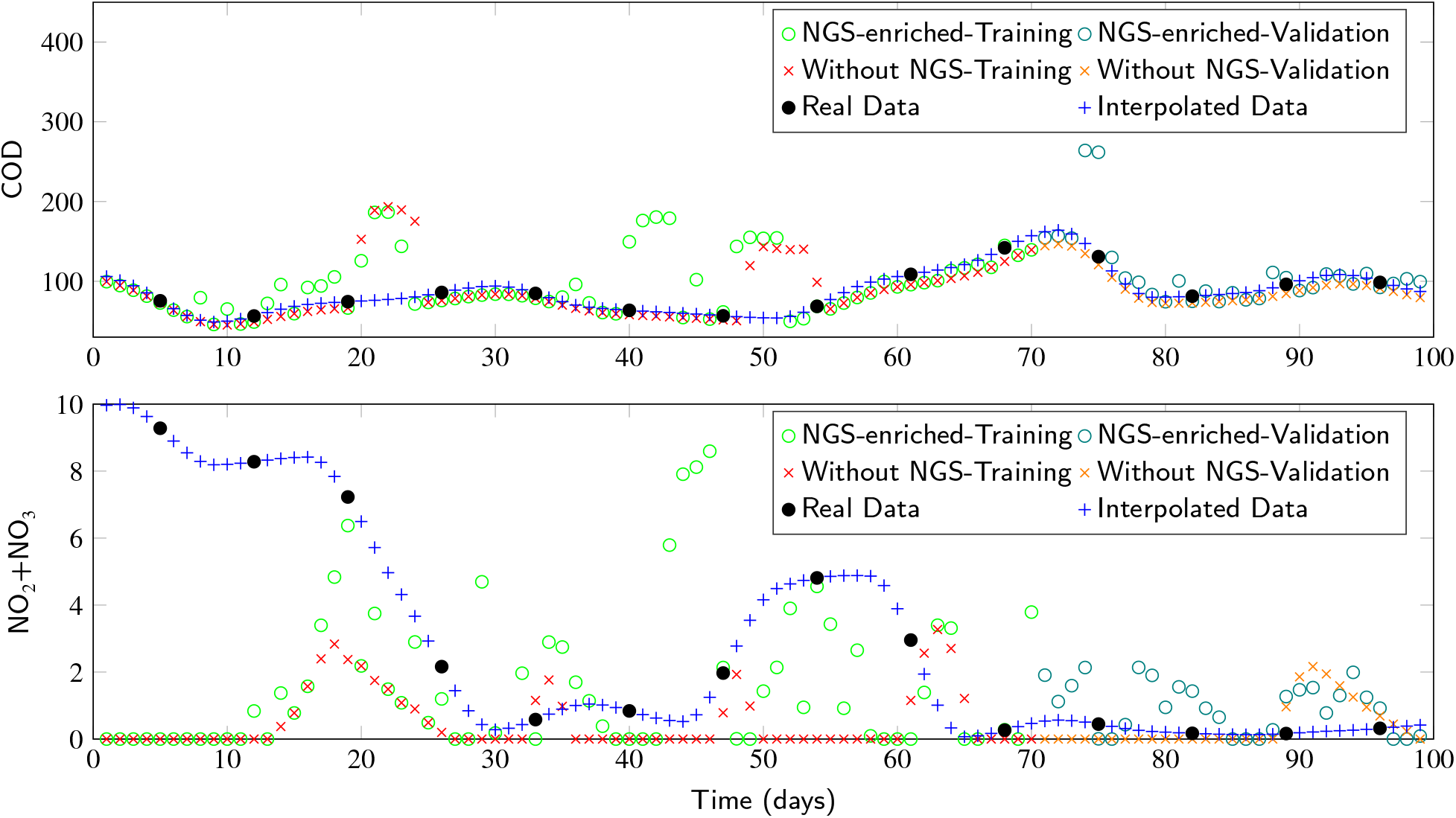
The plot of time-series data of COD and 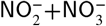 from measurement (shown in blue plusses) and simulations. The simulations give a direct comparison on the use of static mod-ASM1 (red crosses) and on the use of NGS-enriched mod-ASM1 (green circles). In the simulation, the data from day 1 to day 70 correspond to the training data while the data from day 71 to day 100 are the validation data that are used to evaluate the predictive capability of the models.

On the one hand, it can be seen from Figure 4 that the quality of model fitting and prediction for the COD is in a good agreement with the WWTP process data in both cases (with and without the coupling to the NGS data). On the other hand, the fitting and prediction quality of the models to describe the process data of 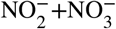 show that the NGS-enriched model can provide a better monitoring signal to the nitrogen production in the effluent than the static one. When we analyze the mean-square-error (MSE) value at the actual sampling time points, the mod-ASM1 model without NGS data gives an MSE value of 15.14 while that with NGS data yields a better MSE value of 11.17. This shows that the NGS data has provided useful information to the model. Albeit the NGS-enriched model requires some time at the start before the signal on 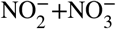 picks up, the correlation between the NGS-enriched model and the real process data is higher than the static model, which gives zero output most of the time.

## 5. Conclusion

In this paper, we presented NGS-enriched coarsegrained modeling of North Water’s Saline Wastewater Treatment Plant that is based on Activated Sludge Model 1. Using a modified ASM1, we enrich it with the NGS data by making two of the parameters time-varying and by coupling them to a subset of the microbial population from NGS data. The enriched model shows a good agreement with the training and validation data on COD, and it can produce a good monitoring signal for the production of nitrogen in the effluent.

A number of improvements can be considered to the current setup. Currently, we used identical parameters (stoichiometric and kinetic) for the two reactor systems of anaerobic and aerobic processes, which can be improved further by taking into account a different set of parameters for each reactor. Another improvement is to increase the frequency of NGS data collection and analysis from four-weekly measurements to daily ones reflecting the fact that the WTTP is highly dynamic, as shown before in Figure 4. Lastly, we can consider the use of other activated sludge models, where the removal of another chemical substance, such as phosphorus, that is abundant in industrial WWTP, can be incorporated.

## Acknowledgement

We thank the Center for Information Technology of the University of Groningen for its support and for providing access to the Peregrine high-performance computing cluster. We also thank A.K. Geurkink for the great discussion. The data used in this paper is taken from North Water’s Wastewater Treatment Plant in Delfzijl. North Water provided the analytical data of the chemical composition of the influent and effluent, and the microbiological samples of the activated sludge compartment of the WWTP are processed by Bioclear earth. Moreover, M A Prawira Negara profoundly expresses gratitude to the Islamic Development Bank (IsDB) for the scholarship funding.

## Nomenclature

*η_g_*: correction factor of *μ_H_* under anoxic conditions
*η_h_*: correction factor for hydrolysis under anoxic conditions
*μ_A_*: specific growth rate for autotrophic biomass (d^−1^)
*μ_H_*: specific growth rate for heterotrophic biomass (d^−1^)
*b_A_*: decay coefficient for autotrophic biomass (d^−1^)
*b_H_*: decay coefficient for heterotrophic biomass (d^−1^)
*COD*: chemical oxygen demand (mg/L)
*f_p_*: fraction of biomass leading to particulate products
*I_XB_*: nitrogen fraction in biomass
*I_XE_*: nitrogen fraction in products from biomass
*k_a_*: Ammonification rate (d^−1^)
*k_h_*: hydrolysis rate constant (d^−1^)
*K_S_*: half-saturation coefficient for readily biodegradable substrate (mg/L)
*K_X_*: half-saturation coefficient for particulate biodegradable substrate (mg/L)
*K_NH_*: half-saturation coefficient for ammonia nitrogen (mg/L)
*K_NO_*: half-saturation coefficient for nitrate and nitrite nitrogen (mg/L)
*K_OA_*: oxygen half-saturation coefficient for autotrophic biomass (mg/L)
*K_OH_*: oxygen half-saturation coefficient for heterotrophic biomass (mg/L)
*S_I_*: soluble inert organic matter (mg/L)
*S_O_*: dissolved oxygen (mg/L)
*S_S_*: readily biodegradable substrate (mg/L)
*S_NH_*: ammonia nitrogen (mg/L)
*S_NO_*: nitrate and nitrite nitrogen (mg/L)
*X_I_*: particulate inert organic matter (mg/L)
*X_S_*: slowly biodegradable substrate (mg/L)
*X_BA_*: active heterotrophic biomass (mg/L)
*X_BH_*: active autotrophic biomass (mg/L)
*Y_A_*: growth yield of autotrophic biomass
*Y_H_*: growth yield of heterotrophic biomass

## Appendix A. Supplementary data

Supplementary data to this article can be found online at https://data.mendeley.com/datasets/gcdckhx3pt/2 (Negara and Geurkink (2022)).

## Footnotes

1 NGS is a massive parallel or deep sequencing that describes a DNA sequencing technology that has revolutionized genomic research (see Behjati and Tarpey (2013)).

2 The plant is also known by its Dutch acronym “ZAWZI”.

